# Nanoscale rheological heterogeneity revealed by Single Particle orientation Tracking (SPoT) of ultrashort carbon nanotubes in brain tissue

**DOI:** 10.64898/2026.05.04.721587

**Authors:** Limeng Ruan, Hanna Manko, Quentin Gresil, Luis A. alemán-Castañeda, Morgane Meras, Finn L. Sebastian, Ivo Calaresu, Benjamin Flavel, Jana Zaumseil, Laurent Groc, Sophie Brasselet, Marc Tondusson, Laurent Cognet

## Abstract

Transport in complex biological tissues is governed by local rheological heterogeneity at the nanoscale, yet probing such environments deep inside living systems remains challenging. Here, we introduce an orientation-sensitive single-particle tracking (SPoT) approach that simultaneously resolves translational and rotational dynamics of individual carbon nanotubes deep within biological tissue. By exploiting the intrinsic dipole-like emission and short-wave infrared luminescence of carbon nanotubes enhanced through the incorporation of quantum color-centers our method enables long-duration tracking with high signal-to-noise ratio in optically dense environments. Crucially, the length of these nanotubes can be precisely shortened down to a few tens of nanometers to adapt to diffusion environmental dimensions, further optimizing the tracking applicability. SPoT of single carbon nanotubes provides access to relative changes in local viscosity, steric constraints, and environmental anisotropy. When applied to the brain extracellular space, SPoT demonstrates that local variations in the translational and rotational diffusion of tracers are heterogeneous and not systematically correlated. By combining trajectory-based signatures of confinement with rotational dynamics, this analysis allows the local contributions of effective viscosity and spatial tortuosity to be disentangled. By enabling combined translational and rotational tracking of nano-emitters over unprecedented depths and timescales, this work establishes a new framework for probing nanoscale transport and rheological heterogeneity in intact biological tissues and more generally in complex diffusive environments.

## Introduction

Understanding the local rheological properties of materials is crucial for a wide range of scientific and industrial applications, from biological tissues to porous media and advanced functional materials. In biological systems, local viscosity and viscoelasticity critically regulate fundamental processes such as molecular diffusion [1–3], intracellular transport [4], mechanotransduction [5, 6], and tissue morphogenesis [7]. At the microscale, viscosity governs the behavior of complex fluids and molecular transport, thereby influencing both physiological function and the emergence of pathological states [3, 8, 9]. Yet, accessing local viscosity in heterogeneous and crowded biological environments remains a major experimental challenge [10]. Conventional rheological approaches, including bulk rheometry, provide ensemble-averaged measurements and lack the spatial resolution required to probe the intrinsic nanoscale heterogeneity of tissues and cells. Recent advances in biomechanical imaging, such as shear wave elastography, Brillouin microscopy, and optical coherence elastography, have enabled non-invasive mapping of mechanical properties in biological tissues, but these techniques predominantly report elastic moduli and provide only indirect or limited access to viscous dissipation [11–14]. Nanoindentation and atomic force microscopy offer submicron mechanical resolution, yet they are generally restricted to surface measurements or ex vivo samples and remain poorly suited for probing dynamic viscosity deep inside living tissue [15–17]. Complementary strategies based on molecular and nanoparticle probes were reported to probe microviscosity in living systems. Viscosity-sensitive fluorescent dyes and molecular rotors have revealed spatially resolved microviscosity maps within cells and organelles, highlighting strong correlations with cellular metabolism and disease states [18–20]. Fluorescence lifetime imaging microscopy (FLIM) has further improved quantitative microviscosity measurements; however, its applicability remains limited by penetration depth, scattering, and spatial resolution in thick tissues [21]. As a result, high-resolution, non-invasive approaches capable of probing local viscosity deep within complex biological environments remain fundamentally lacking.

Among such environments, the brain extracellular space (ECS) is of particular interest. In this key compartment for brain function, molecular transport is governed by a subtle interplay between geometric constraints, molecular crowding, charge, pH, molecular shape and local viscosity, all of which strongly modulate diffusion and signaling in the central nervous system [22, 23]. Single-particle tracking (SPT) has transformed the study of nanoscale transport in biological systems, and translational tracking of diffusing probes has provided valuable insights into ECS tortuosity and hindered diffusion [24–31]. However, translational motion alone is insufficient to disentangle viscosity from steric and structural constraints. The rotational dynamics of anisotropic probes provide a complementary readout that is highly sensitive to the immediate nanoenvironment, including viscous drag, steric hindrance, molecular interactions, and nanoscale anisotropy. Joint analysis of rotational dynamics and translational trajectory features should therefore enable viscous and structural contributions to the local nanoenvironment to be distinguished, revealing properties that remain inaccessible through positional tracking alone. The timescales of molecular reorientation may greatly vary depending on the size, anisotropy of the nanotracers to be studied in a specific environment spanning nanoseconds to millisecond and higher. Fluorescence anisotropy which measures fast rotational dynamics lacks spatial resolution and sensitivity to reach the single molecule regime. On the other hand, a number of powerful methods have been recently developped to measure the 3D orientation of single emitters, including polarization splitting methods and point spread function (PSF) engineering methods (called ‘Dipole Spread Function’ DSF engineering). These methods have been successfully applied to the study of fixed systems and snapshot measurements in well-controlled model environments, including supported lipid bilayers, two-dimensional crystals, and cell membranes. However, none of these methods could be applied to tracking the orientation of single molecules diffusing in complex environments. This limitation is primarily due to the lack of adequate probes that require orientational dynamics to be resolved into diverse environments and low photobleaching for extended measurements [32–34]. Scattering-based approaches, including differential interference contrast (DIC), defocused dark-field microscopy and cylindrical-polarization-based interferometric scattering microscopy (cypiSCAT), have also been applied to rotational tracking but are generally confined to surface-bound probes or thin samples [35–38]. To date, orientational tracking with long trajectories in deep, optically dense tissues has remained out of reach.

Here, we broke these limitations by introducing an approach to perform single particle orientation tracking (SPoT) based on dipole spread functions (DSF) engineering. This method simultaneously resolves translational and rotational dynamics at tens of milliseconds timescales, deep within biological tissue over extended recording times, and with nanoscale spatial precision. Our method exploits short-wave infrared (SWIR)–emitting carbon nanotubes (CNTs) as uniquely suited one-dimensional probes, combining exceptional photostability, strong dipolar emission, and minimal scattering for deep tissue imaging [39,40]. Importantly, by adapting CNT lengths to the environmental conditions, we demonstrate long-term, high-precision tracking in complex biological environments. Building on previous work using SWIR-emitting CNTs to probe molecular diffusion in the brain ECS [24–26, 31, 41–43], SPoT extends their capabilities to orientational tracking.

For the first time, we demonstrate simultaneous long-duration tracking of both position and orientation of individual nanoparticles deep inside living biological tissue. By combining advanced polarization microscopy, dynamic super-localization of SWIR 1D nano-emitters, and deep-learning–based analysis, our approach provides access to local nanoscale signatures of viscosity heterogeneity and environmental anisotropy that are inaccessible to existing techniques. Applying this strategy to the brain ECS, we reveal how extracellular matrix integrity, molecular crowding, and heterogeneous local environments modulate both translational and rotational dynamics. These findings establish a powerful framework for probing nanoscale transport and rheological heterogeneity in intact biological tissues.

## Results and discussion

CNTs are one-dimensional nano-emitters in the SWIR domain which cover the NIR-II window [40]. They exhibit high brightness and photostability. Their nanoscale dimensions enable them to diffuse more effectively through complex and confined structures, making them excellent probes for nanoscale bioimaging in deep tissue [24, 44–50]. Beyond their optical advantages, the anisotropic one-dimensional geometry of CNTs provides unique information during diffusion processes. Indeed, length influence rotational diffusion. Precise control over CNT lengths provides thus a unique way to tailor diffusion dynamics to diverse environments. Provided that the nanotubes are sufficiently bright, this capability enables simultaneous measurements of translational and rotational dynamics by single-molecule microscopy. To achieve this, we employ CNTs functionalized with quantum defects, which locally trap excitons and create emissive color centers (CCs) along the nanotube backbone [51–54]. Specifically, we synthesized bright, biocompatible long and ultrashort CNTs functionalized with oxygen CCs emitting at 1120 nm. These nanotubes can be excited either at 845 nm via a phonon sideband or at 985 nm through the first excitonic transition [43]. These are referred to hereafter as CCNTs and ultrashort CCNTs (uCCNTs), with respective lengths of 550 nm and 50 nm (Figure S1). Importantly, within this length range, the limited spatial resolution of optical microscopes prevents the direct visualization of CCNT orientation. As a result, CCNTs systematically appear as diffraction-limited punctual emitters in recorded images.

Due to their characteristic geometry, CCNTs behave as emitting dipoles lying along the nanotube longitudinal axis, whose direction can be monitored to retrieve their orientation. To avoid undefined values of *ϕ* at the poles (*θ* ∼ 0^◦^) in spherical coordinate system the orientation can be described using a unit vector

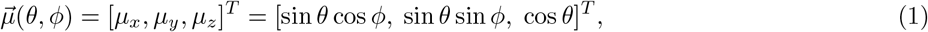

where z is the optical axis, *θ* is the polar angle and *ϕ* is the azimuth angle (Figure 1(a)).

**Fig. 1.**
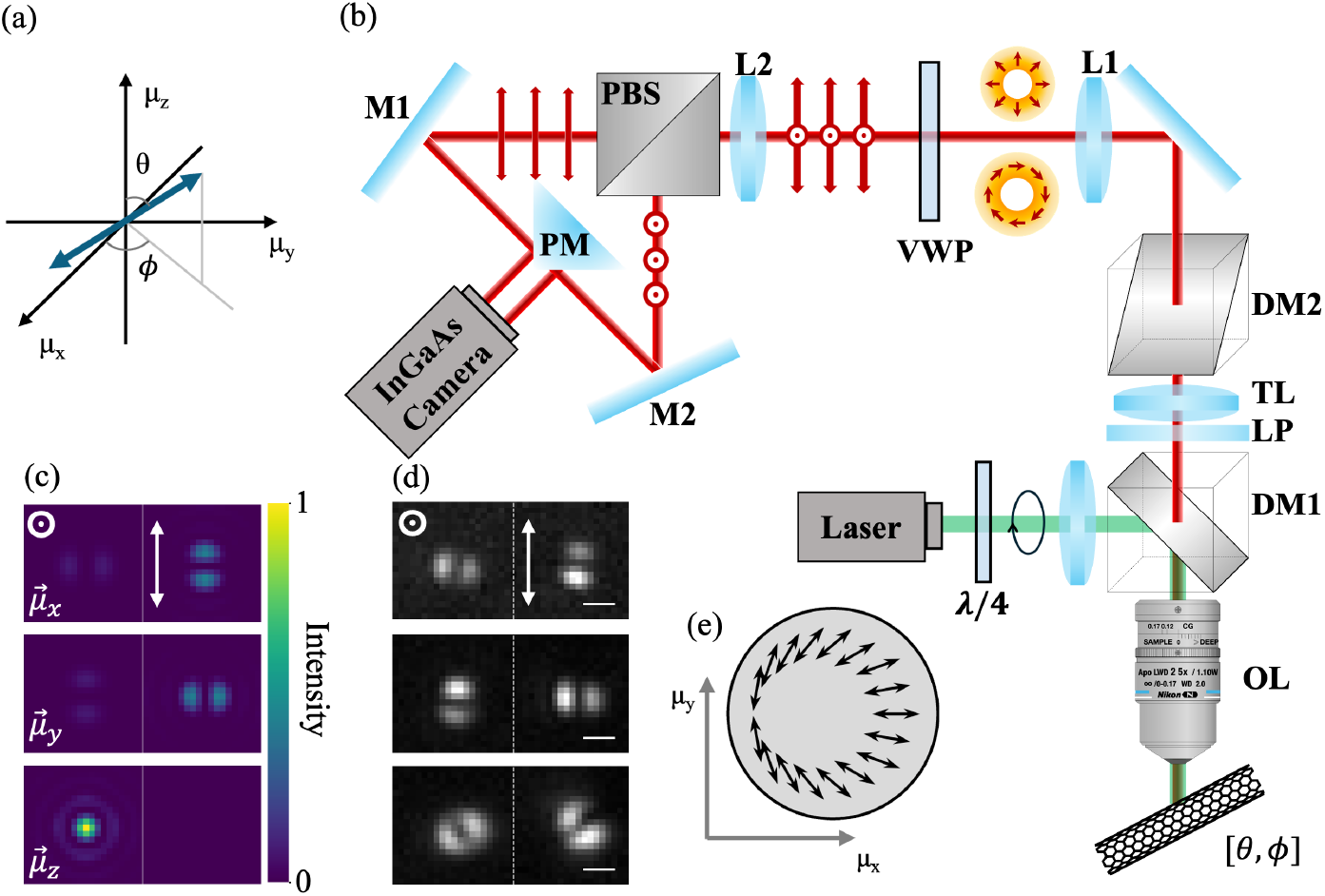
(a) The orientation of the CCNT emission dipole is described using a unit vector 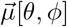. (b) raPol microscope setup schematic. The components are described in the section methods. (c) Simulated DSFs generated by the raPol microscope when dipoles along *µ*_*x*_, *µ*_*y*_, *µ*_*z*_ axis, respectively. Color bar: intensity. (d) Examples of measured DSFs by raPol microscope. Scale bar: 1 *µ*m. Left: X-Radial DSF. Right: Y-Azimuthal DSF. (e) Schematics of a VWP. Arrows represent the VWP’s fast axis.

In order to enable real-time measurement of the 3D orientation (*θ, ϕ*) and 2D position (x,y) of individual CCNTs along diffusive trajectories, we developed a radially and azimuthally polarized (raPol) microscope operating in the SWIR range [55]. The method relies on a single-molecule epi-fluorescence wide-field microscope, where the sample is excited using a circularly polarized laser and the emitted fluorescence is analyzed through polarization engineering at the objective’s back focal plane (BFP) (Figure 1(b)). More precisely, at the BFP which is accessible using an imaging relay system, a vortex half-wave plate (VWP) converts radially and azimuthally polarized components of the fluorescence into two orthogonal linear polarization states. These components are subsequently separated by a polarizing beam splitter, producing two spatially distinct images that are projected side-by-side onto a InGaAs camera allowing detecting single CCNTs emitting in the SWIR [54]. The resulting images encode the orientation information of the emitting dipole. For emitters aligned along the optical axis, the fluorescence is confined to a single polarization channel (x-Radial image, Figure 1(c)). In contrast, emitters oriented within the transverse plane produce DSFs consisting of two lobes in the y-Azimuthal image. In the azimuthal polarization channel, the line connecting the lobe centers directly corresponds to the in-plane orientation angle *ϕ*. For random orientations, both channels contain information (Figure 1(d)) and by analyzing the relative intensity distribution and spatial structure of these DSFs, the 3D orientation and in plane super-localization of CCNTs can be quantitatively determined.

In practice, due to inherent aberrations of the optical assembly, experimental DSFs differ from the exact symmetric double-Gaussian-like shape expected from theory. To retrieve orientational parameters in an accurate way from experimental DSFs, it is nevertheless crucial to account for all possible effects at the origin of DSF-deformations i.e. phase and polarization distorsions present in the emission path of the microscope. To estimate such distorsions in the relay pupil plane of the microscope objective, we use a vectorial phase retrieval method [56] which provides the birefringent distribution at the pupil plane (BDPP). The BDPP can be modeled by a spatially varying Jones matrix, which contains the aberrations and polarization distortions of the setup. In a raPol microscope without optical aberrations, the effective Jones matrix of VWP is given by [57]

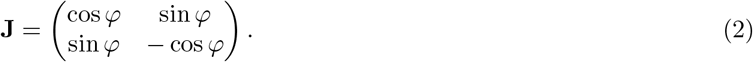

To express the matrix elements relative to our experimental setup, we need to take into account all possible states of polarization. Because generating samples with all possible 3D orientations is challenging, a solution is to use nanoparticles which emit unpolarized fluorescence and to generate both a phase and polarization diversity in the emission path by introducing retardation plates. More precisely, we use 100 nm polymeric nanoparticles embedded with gold nanoclusters (GNPs) [58] which emit unpolarized SWIR fluorescence light. These nanoparticles were immobilized on a polylysine coated coverslip for imaging.

By combining polarization and phase diversities (Figure 2(a)), the point spread functions (PSFs) generated from the light emitted by GNPs (Figure 2(b)) in the field of view (2×40×40 *µ*m) are then processed by a nonlinear optimization algorithm to retrieve the experimental BDPP (Figure 2(d)). For this, we averaged 20 images acquired at 100 ms integration time. When comparing simulated and experimental BDPPs (Figure 2(c,d)), the experimental system still suffers from slight misalignement evidenced by the shift between the BDPP centers. As well, despite careful optical design, some polarization distorsions in the microscope are visible from the experimental BDPP. Figure 2(e) shows simulated DSFs for dipoles at various orientations, based on the experimental BDPP which accounts for optical aberrations of the microscope, including polarization aberrations. The simulated DSFs are in close agreement with the experimental observations (Figure 1(d)) validating the BDPP retrieval. We next compared the theoretical best achievable localization and orientation precision, as predicted by the Cramér–Rao lower bound (CRLB; Figure S2), using simulated and experimentally measured BDPPs (Figure 2(c,d)) [59, 60]. Although experimental aberrations slightly increase the predicted uncertainties, they remain within a practically useful range.

**Fig. 2.**
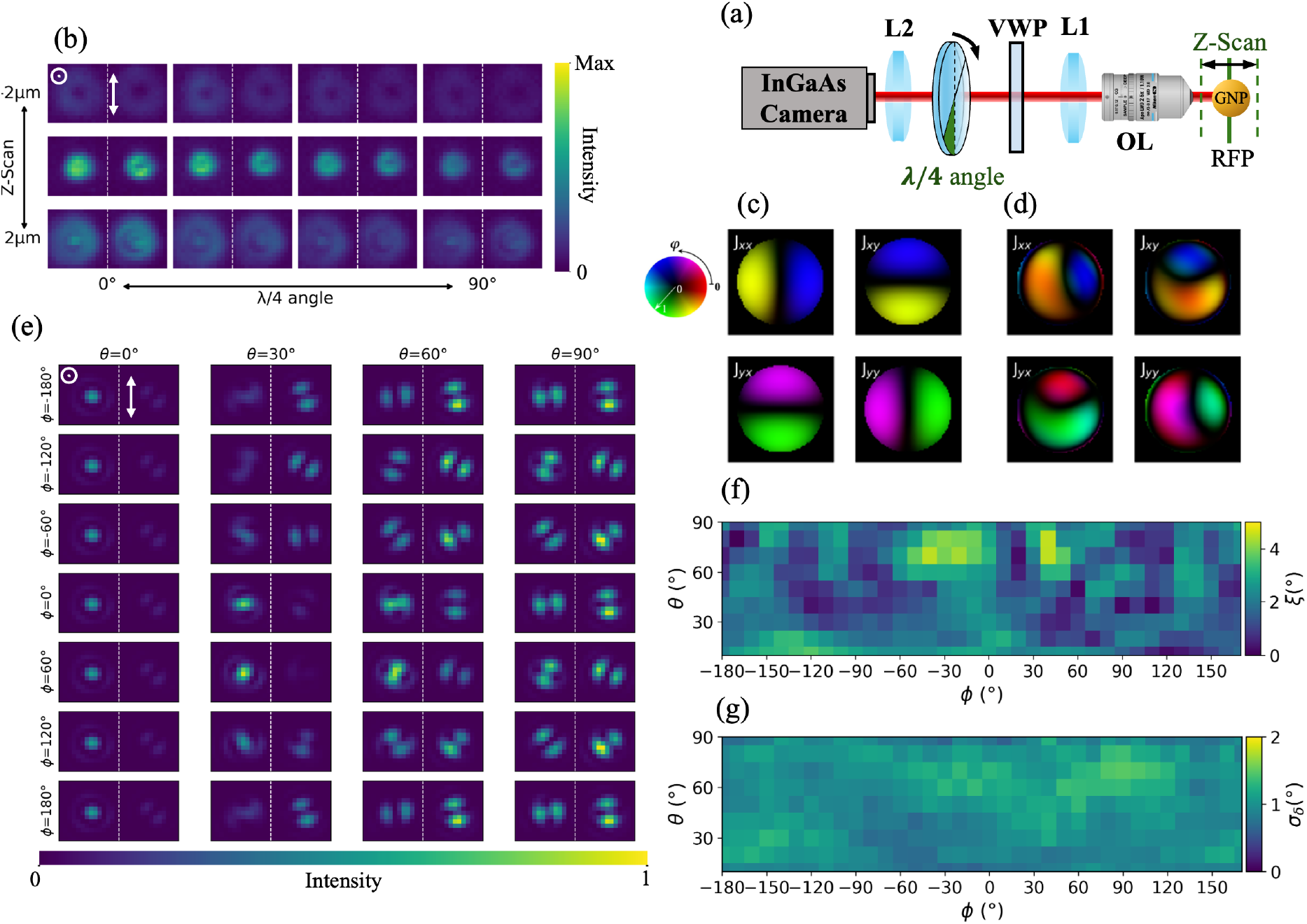
(a) Schematic of polarization projection diversity (λ/4 angle) and phase diversity (Z-Scan) for a GNP. (b) Measured phase and polarization diversity stacks. Color bar: intensity. (c) The retrieved elements of the Jones matrix for the BDPP from numerical experiment without aberrations. (d) The retrieved elements of the Jones matrix for the BDPP from experimental data. (e) The DSFs generated by all orientation dipoles simulated using the retrieved BDPP in (d). Color bar: intensity. Left: X-radial PSFs. Right: Y-azimuthal PSFs. (f,g) Deep-SMOLM estimation accuracy (f) and precision (g). (f) Orientation accuracy (*ξ*). (g) Mean angular standard deviation *σ*_*δ*_.

Introducing the extracted Jones Matrix of the BDPP in a deep-learning algorithm (Deep-SMOLM), allows then to analyze and obtain the 2D localization and 3D orientation of single CCNTs from the raPol microscope [61]. In order to train the algorithm, we numerically generated 2000 images based on the experimental BDPP. For each training image, we randomly set the number of dipole emitters between 5 and 10, each of them having random 2D localization and 3D orientation. Furthermore, during training we randomly set the focal imaging position *z* ∈ [−500 nm, 500 nm], allowing the network to learn the effects of defocus and thus produce reliable reconstructions even when particles are slightly out of focus. We also set the emitter intensity following a Poisson distribution (rate parameter = 4) with additive Gaussian noise (*µ*_*G*_ = 0, *σ* = 2), followed by linear gain, offset correction, and thresholding at 50, and is finally scaled by a photon number factor of 500. Furthermore, a uniform background of 20 photons per pixel is added to all pixels. Finally, Poisson noise is added to the images.

We first aimed at quantifying the accuracy and precision of our approach to retrieve CCNT orientations and localizations. For this, we generated simulated images of individual emitters with random fixed positions, intensities, and orientations. For each orientation, we analyzed 100 different images with 500 photons. The (non-negative) polar and azimuthal angular distances between the estimated orientation 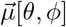 retrieved from Deep-SMOLM and the ground truth orientation 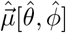 were calculated to assess the orientation accuracy (*ξ*)

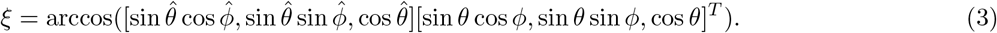

On the other hand, the orientational precision *σ*_*δ*_ is obtained from the estimated orientation 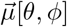, which represents the angle of an equivalent circular uncertainty region on the unit sphere. It provides a single scalar measure of orientation precision by combining the estimation uncertainties in both the polar and azimuthal directions (see Supporting Information) [61, 62],

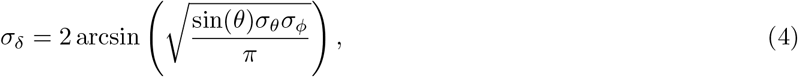

where *σ*_*θ*_ and *σ*_*ϕ*_ are the standard deviation of measuring *θ* and *ϕ*. For all accessible values of *θ* and *ϕ*, we obtain an accuracy (Figure 2 (f)) which is better than 5^◦^, and a precision (Figure 2 (g)) better than 2^◦^. In addition, the localization precision are below 8 nm (Figure S3). We estimate that such performances are consistent with CRLB predictions making them promising enough to allow experimental determination of single CCNT orientation and localization in one shot.

In a first experimental test, we analyzed a sample of (6,5) chirality uCCNTs coated with deoxycholate (DOC) (length ≈50 nm), exited at 845 nm. These uCCNTs were immobilized on a glass coverslip by spin-coating them onto a polyvinylpyrrolidone layer. This configuration was used to validate the performance of our system with uCCNTs predominantly oriented within the (xy) plane, while allowing for a certain degree of out-of-plane misalignment induced by the polymer layer. For this test, we have averaged 50 consecutive frames (100 ms per frame) before analyzing uCCNTs orientations and localizations as shown in Figure 3(a). Examples of experimental DSFs from CCNTs immobilized on similar surfaces are shown in Figure S1 and exhibit patterns identical to those observed for uCCNTs. Figure 3(c) shows all the merged results and we found that *ϕ* displays a uniform distribution, while most of *θ* values are confined between 60−90^◦^. This indicates that as expected after spincoating, most 1D uCCNTs are close to parallel to the surface of the coverslip yet with a small narrow out-of-plane distribution due to the use of the polymer layer. We next repeated the same experiment, where uCCNTs were embedded and immobilized in 5% agarose gels [63]. In this case, uCCNTs orientations are expected to be randomly distributed in 3D which is observed by our analysis Figure 3(c). Indeed, in this situation, *θ* spans a much broader range than for spincoated nanotubes. Interestingly, we noticed that uCCNTs oriented close to the optical axis of the microscope 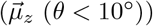 could not be observed. We attribute this effect to the fact that such nanotubes could not be excited efficiently due to the lack of axial polarization component in the excitation beam (which was set to circular polarization). Finally, we analyzed uCCNTs embedded in 5% agarose gel at depths, ranging from 0 to 200 *µ*m. Over this range, we observe no significant degradation in either localization or orientation precision with increasing imaging depth (Figure S4). Consequently, within the investigated depth range in such samples, imaging depth has only a minor effect on the detected photon budget and the measured DSFs.

**Fig. 3.**
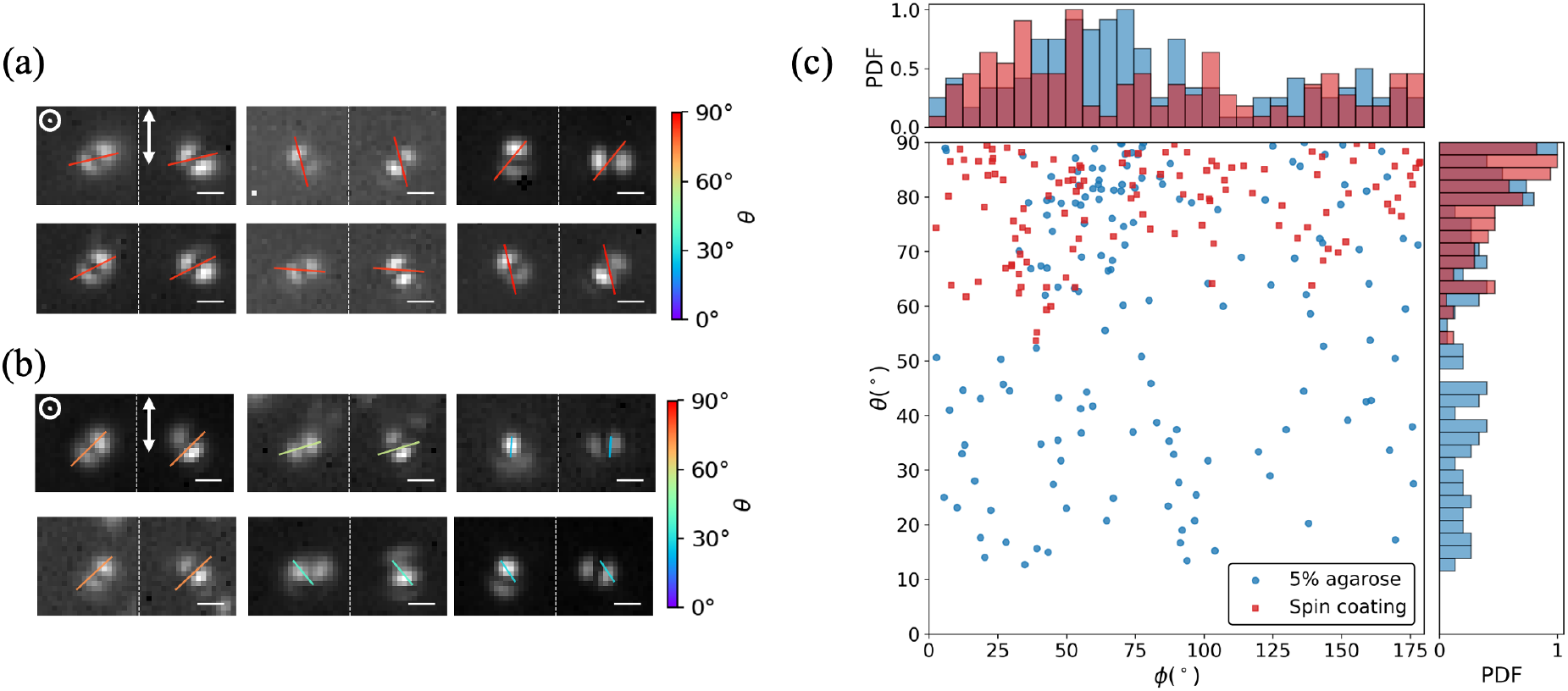
The orientations of uCCNTs are fixed by spin-coating and 5% agarose. (a, b) uCCNTs raPol images and their orientation results. The line indicates the azimuthal angle *ϕ* of each uCCNTs, and its length and color corresponds to the polar angle *θ*. The center of the line corresponds to the spatial coordinates of the emitter. Color bar: *θ*. Left: X-radial DSFs. Right: Y-azimuthal DSFs. Scale bar: 1 *µ*m. (c) All orientation result and probability density function (PDF) of *θ* and *ϕ* of spin-coating (red) and 5% agarose (blue). For a more compact representation, we display *ϕ* over the range [0^◦^, 180^◦^], with negative values wrapped into this interval.

Such performance where single shot images allow 3D orientational and 2D localization determination of single CCNTs opens the possibility for performing dynamic studies based on single particle tracking experiments. Our next objective was thus to compare rotational diffusion derived from 3D orientation with translational diffusion obtained from 2D positional trajectories.

We first conducted dynamic experiments in which CCNTs were dispersed and freely diffused in glycerol/water mixtures at varying concentrations (90%, 80%, and 70%, w/w). Examples of measured DSFs are presented in Figure S5. In such media CCNT show unconstrained translational and rotational motion. Because the rotational diffusion of 1D objects is strongly dependent on their length and on the viscosity of the surrounding medium (hence the use of high glycerol concentrations), using 33 ms exposure times, SPoT enabled us to resolve orientational diffusion at the single-nanoparticle level when using the long CCNTs sample having an average length of approximately 550 nm. In such unconstrained environments, much shorter CCNTs would rotate too rapidly for their orientational state to be accurately resolved and tracked at the integration times required to detect single CCNTs.

The trajectories represented on Figure 4(a) allow us to analyze the diffusion characteristics of the CCNTs in the three different glycerol/water mixtures. For this we approximate CCNTs by ellipsoids which can rotate independently around each of its three axes [64], The long axis is the molecule’s orientation and the two short axes are equal in length, much smaller than the long axis. Over a short time interval Δ*t*, the angular displacements around these axes are denoted as (*δφ*_*a*_, *δφ*_*b*_, *δφ*_*c*_). Since rotation around the long axis (*δφ*_*a*_) does not alter the molecule’s orientation, we set *δφ*_*a*_ = 0^◦^, and the rotational diffusion around the two short axes is characterized by a single diffusion coefficient, *D*_*r*_. Its dependence is based on the shape of the molecule and dynamic viscosity of the medium, *η*. For CCNTs in the slender-body approximation we can write

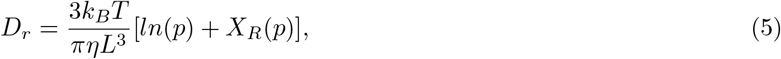

where *k*_*B*_ is the Boltzmann constant, *T* is the absolute temperature, L is the length, d is the diameter, *p* = *L/d* is the aspect ratio, and *X*_*R*_(*p*) is a finite-length correction term [65].

**Fig. 4.**
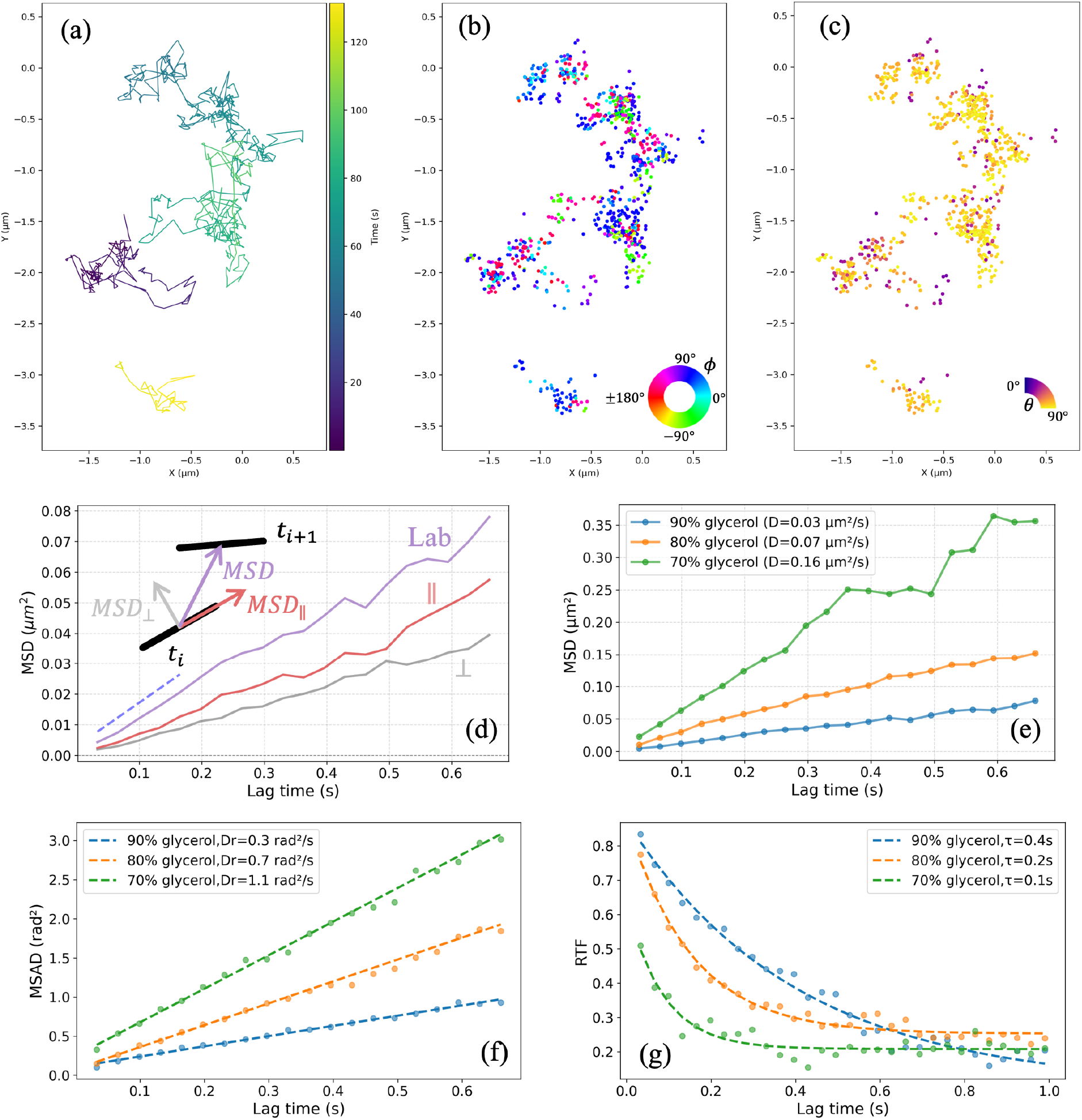
The trajectories and analyzed results of CCNTs in glycerol. (a–c) A trajectory of CCNTs in 90% glycerol. Color bar: (a) time, (b) azimuthal angle *ϕ*, (c) polar angle *θ*. (d) MSDs of CCNTs with *D*_‖_ = 0.04 *µ*m^2^*/*s and *D*_⊥_ = 0.03 *µ*m^2^*/*s) for 90% glycerol. Blue dashed line: the slope is 2(*D*_‖_ + *D*_⊥_). MSDs in the lab frame for 90%, 80%, and 70% glycerol. (f) MSADs, and (g) RTF for CCNTs in 90%, 80%, and 70% glycerol.

For Brownian diffusion, the angular displacements in the nanotube body frame (*δφ*_*b*_ and *δφ*_*c*_) follow a normal distribution: *δφ*_*b*_, *δφ*_*c*_ ∼ *N* (0, 2*D*_*r*_Δ*t*), where *N* represents a normal distribution. The total body frame displacement is then calculated by summing the displacements over each time step.

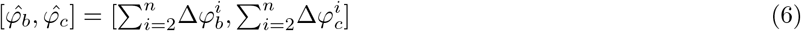

where i is *i*_*th*_ time step. The mean-squared angular displacement (MSAD) can then be expressed as

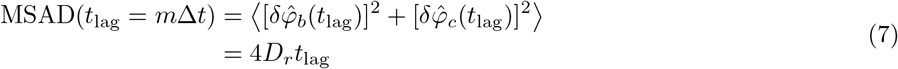

This indicates that one can obtain *D*_*r*_ from the local slope of the MSAD.

For an anisotropic particle, translational motion can be decomposed into components parallel and perpendicular to its orientation characterized by two independent diffusion coefficient *D*_‖_ and *D*_⊥_, respectively. Having access to the orientation of a single CCNT thus allows to express two mean-square displacements (MSDs) in the CCNT body frame 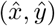,

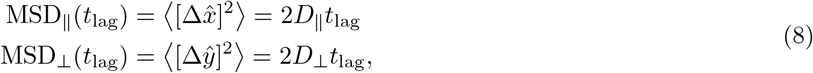

which can also be used to obtain in the laboratory frame where the particule coordinates are (x, y) one can also defined the mean-square displacement (MSD) in the laboratory frame (*x, y*)

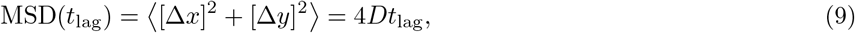

here where

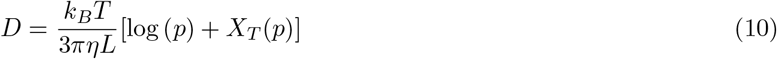

is diffusion coefficient and *X*_*T*_ (*p*) is also a finite-length correction term [65].

Interestingly, the MSDs calculated in the lab frame and in the CCNT body frame are linked by the following relation [66]:

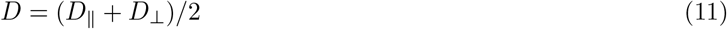

By incorporating orientation information, our approach enables MSD analysis in the nanotube frame, referred in this work as the body-frame, allowing the anisotropic translational diffusion of individual nanotubes to be quantified. In addition, we introduced the MSAD, which provides direct access to rotational diffusion.

Having the CCNT orientation, we can also calculate the reorientational timecorrelation function (RTF) [67,68] which allows to determine the rotational correlation time (*τ*) of the nanotube movement,

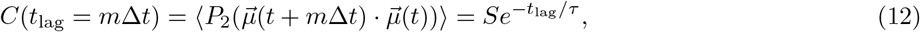

where S is constant. In practice *τ* is the characteristic timescale over which molecular orientations lose memory of their initial direction and *P*_2_ is the second-order Legendre polynomial.

Figure 4(a) displays the Brownian trajectories of a CCNT in 90% glycerol recorded by SPoT. Accordingly, displaying *θ* and *ϕ* at each position in the trajectory (Figure 4(b-c)) indicates that the CCNT experiences random orientations along the trajectory. A total of 16 such trajectories were analyzed, and the time-ensemble-averaged MSD was further calculated [69] and displayed in Figure 4(d). As expected, the MSD of CCNTs subject to Brownian motion calculated in the lab frame is linear, and a linear fit yields a diffusion coefficient of *D* = 0.03 *µ*m^2^*/*s. In the CCNT body frame, the diffusion along the long axis (*D*_‖_) is faster than that along the short axis (*D*_⊥_), and the diffusion coefficient satisfies the expected relation (11). For rotational diffusion, a linear fit to the MSAD gives a rotational diffusion coefficient of *D*_*r*_ = 0.3 rad^2^*/*s (Figure 4(f)), which matches the expected value of 0.5 rad^2^*/*s assuming CCNTs of 550 nm length, 2.3 nm diameter and based on a value of *η*_90%_ = 0.25 Pa·s for 90% glycerol determined by SPT of calibrated spherical particles (*R* = 12 nm,*D* = 0, 07 *µ*m^2^*/*s)). Finally, we computed the RTF (Figure 4(g)) and we find a *τ* = 0.4 s. Since glycerol is an isotropic liquid, the rotational correlation time satisfies *τ* = 1*/*6*D*_*r*_.

We next compared the results obtained for the different glycerol water concentration (90%, 80%, and 70%, w/w) and displayed the results in Figure 4(e-g) and Figure S6. Qualitatively, we find that as the viscosity increases, both the translational diffusion and the rotational diffusion of single CCNTs slow down, while *τ* increases accordingly.

Quantitatively, considering Equation 10 and 5, the difffusion coefficients depend on both the CCNTs size (*L, d*) and the environmental viscosity (*η*). The localization precisions (in x and y) during tracking are *σ*_90%_ = 36 nm, *σ*_80%_ = 61 nm, *σ*_70%_ = 99 nm as retrieved from the intercept at zero time delay [70]. As we used identical CCNT solution, the influence of CCNTs size is effectively averaged out. We thus expect that both translational and rotational diffusion coefficients should follow the same scaling relation *D, D*_*r*_ ∝ 1*/η* and for the same type of particle, the ratios of diffusion coefficients in environments with different viscosity are theoretically equivalent. We determined the viscosity scaling using the same calibrated spherical particles as previously described and obtained the following ratios between the 3 water:glycerol solutions

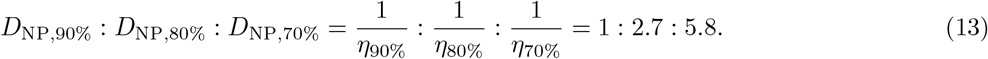

Such values can now be quantitatively compared to the translational diffusion coefficients obtained with CCNTs (Figure 4(e,f)),

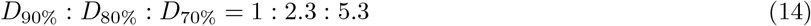

and

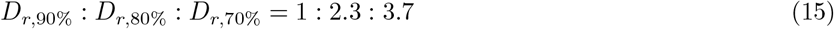

The translational diffusion coefficient ratio obtained with 1D CCNTs closely matches those obtained by the spherical NPs validating the experimental recordings. Concerning the rotational diffusion coefficient measured for CCNTs, excellent agreement is observed for 90% and 80% glycerol. In contrast, for 70% glycerol, the experimentally measured rotational diffusion coefficient is lower than the theoretically expected value. We attribute this observation to the fact that, although the exposure time is sufficiently short for accurate translational tracking, the rotational diffusion of CCNTs in 70% glycerol is too rapid to permit reliable orientation determination over the same acquisition time. As a practical criterion, the characteristic reorientation time should be at least three times longer than the exposure time, ensuring that the particle orientation remains approximately constant during image acquisition. This condition is no longer satisfied in 70% glycerol, thereby defining the upper limit of the rotational diffusion coefficient that can be reliably measured with our setup.

While the glycerol experiments allow unrestricted motion in a homogeneous viscous environment, most biological media impose structural constraints and often exhibit spatial heterogeneities in viscosity. To replicate steric hindrance in a uniform viscosity environment, we next examined the diffusion of CCNTs within 2% aqueous agarose gels. This system provides a controlled porous matrix that selectively restricts molecular motion, thereby mimicking the effects of spatial confinement. Figure 5(a-d) show a CCNT trajectory obtained in 2% agarose gels where translational and angular positions of the CCNT are color coded. As in any SPT experiment based on 2D localization, particles that remain within the focal volume are more likely to be detected over many consecutive frames and therefore yield longer trajectories, as illustrated in Figure 5. Conversely, particles that move out of the imaging volume generate shorter and less informative trajectories. This defocalization, or trajectory-selection, bias is a well-recognized limitation of 2D SPT [71]. Qualitatively one unambiguously observe that in such environments, rotational diffusion is partially hindered by the agarose network (Figure 5(d)). This situation has been extensively studied previously for long nanotube which length could be optically resolved (typically lengths > 1 *µ*m) and which could be either considered as rigid rods like here, and semi-flexible filaments when nanotubes are significantly longer [72, 73].

**Fig. 5.**
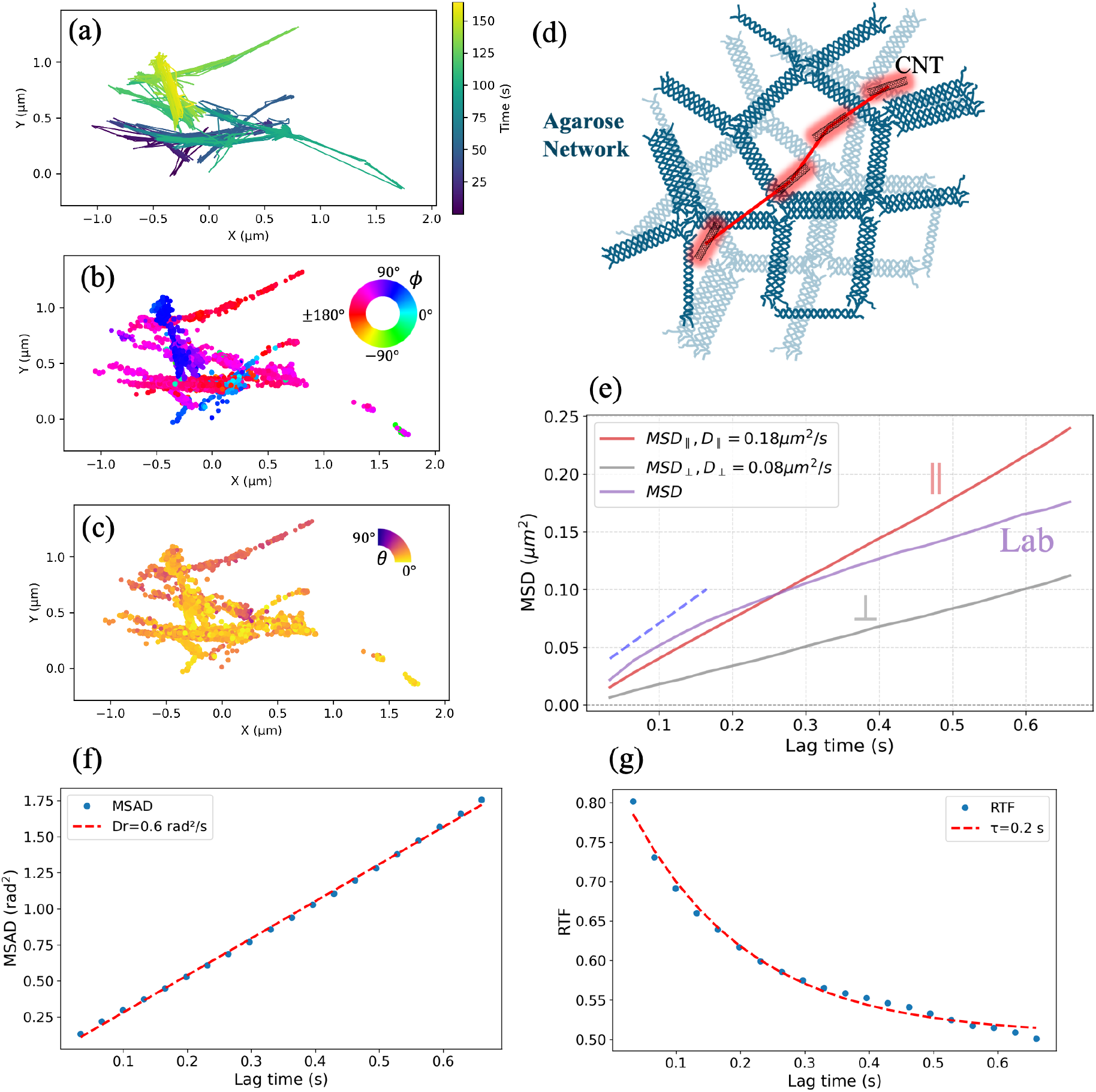
(a–c) Trajectories of CCNTs in 2% agarose. Color bar: (a) time, (b) azimuthal angle *ϕ*, (c) polar angle *θ*. (d) Schematic of an CCNT moving within the 2% agarose network. (e) MSDs of CCNTs in 2% agarose gel. Blue dashed line: the slope is 2(*D*_‖_ + *D*_⊥_). (f,g) MSAD (f) and RTF (g) for CCNTs in 2%agarose. 9 trajectories were analyzed.

Accordingly, we observe owing to the rotational tracking of CCNTs that cannot be resolved optically, that in 2% agarose gel, such 550 nm long CCNTs can not rotate freely; instead, they undergo reciprocating motion along their longitudinal axis until reorientation occurs in the next agarose pocket (Figure 5(a-c)). We next calculate the MSD in the laboratory frame and observe a sublinear dependence, whereas the MSDs calculated in the body frame exhibit a linear behavior. This behavior, previously reported [24], arises from environmental constraints that induce anomalous diffusion over the timescales relevant to the exploration of local gel regions. Analysis of rotational diffusion based on MSAD and RTF yields *D*_*r*_ equal to 0.6 rad^2^*/*s and *τ* equal to 0.2s (Figure 5(f,g)).

The glycerol and agarose experiments establish an operational framework for distinguishing viscous and structural contributions. In homogeneous glycerol solutions, where structural confinement is absent, correlated changes in *D* and *D*_*r*_ primarily reflect changes in viscous resistance. In agarose, confinement is independently identified from the subdiffusive laboratory-frame MSD, trajectory geometry, preferential motion along the nanotube axis, and restricted reorientation. Thus, effective viscosity and structural confinement are inferred from the combined translational and rotational observables, rather than from *D*_*r*_ alone.

We finally demonstrate our methodology to a biological application, namely the study of molecular diffusion in the extracellular space (ECS) in mouse organotypic brain slices. As mentioned in the introduction of this article, translational SPT opened the route toward a finer understanding of molecular diffusion hallmarks in the tortuous ECS but our understanding of the respective impact of viscosity and ECS structure remains to be elucidated [74] and we hypothesize that rotational diffusion tracking will provide novel insights. The ECS is indeed a network of interconnected microdomains bounded by neuronal membranes and filled with interstitial fluid and the extracellular matrix (ECM). In such a system, the diffusion of one-dimensional objects is both highly constrained and heterogeneous, owing to spatial confinement and local variations in effective viscosity [24, 25, 28]. Here, the term effective local viscosity refers to the non-geometric resistance to probe motion arising from the interstitial fluid, molecular crowding, and transient interactions, whereas geometric effects of the ECM are assessed separately from trajectory morphology and orientation constraints. To probe this diversity of behaviors while retaining sensitivity to rotational diffusion, we used uCCNTs (length ≈ 50 nm) instead of the long CCNTs as long nanotubes would not rotate efficiently in the crowded brain ECS. More precisely, we used biocompatible uCCNTs coated with phospholipid–PEG. Moreover, their emission in the biological transparency window reduces tissue scattering and absorption, enabling imaging deep within biological tissues. To assess the effect of imaging depth on the performance of the SPoT method in a complex biological environment, we first imaged immobilized uCCNTs through acute mice brain tissue slices of varying thickness using 100 ms exposure time (Figure S7). As expected, the orientation and localization of uCCNTs could be determined reliably with excellent precision at imaging depths of up to 80 *µ*m (Table 1). At depth approaching 100 *µ*m, however, orientation determination becomes considerably more challenging, primarily due to the reduced photon budget.

**Table 1:**
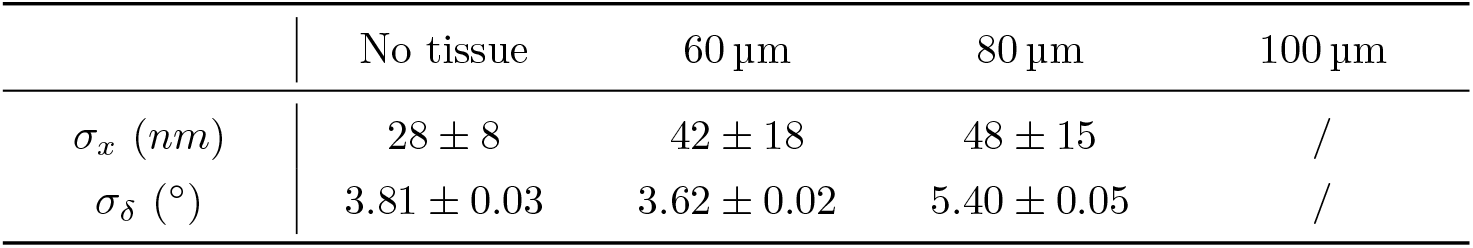
Localization and orientation precision at different imaging depths.

| | No tissue | $60\text{ }\mu\text{m}$ | $80\text{ }\mu\text{m}$ | $100\text{ }\mu\text{m}$ |
| --- | --- | --- | --- | --- |
| $\sigma_x$ (nm) | $28 \pm 8$ | $42 \pm 18$ | $48 \pm 15$ | / |
| $\sigma_\delta$ ($^\circ$) | $3.81 \pm 0.03$ | $3.62 \pm 0.02$ | $5.40 \pm 0.05$ | / |

For diffusion experiments, we next used organotypic slices which allow several aspects of structural connectivity of the original tissue to be preserved [75]. Incubation with uCCNTs-containing medium enables widespread probe penetration and yields a substantially larger number of trajectories, facilitating robust statistical analysis (Figure 6(a)). Markedly, during long-term high-resolution imaging, biological samples often undergo mechanical drift. Therefore, drift correction was applied to compensate for temporal positional shifts.

**Fig. 6.**
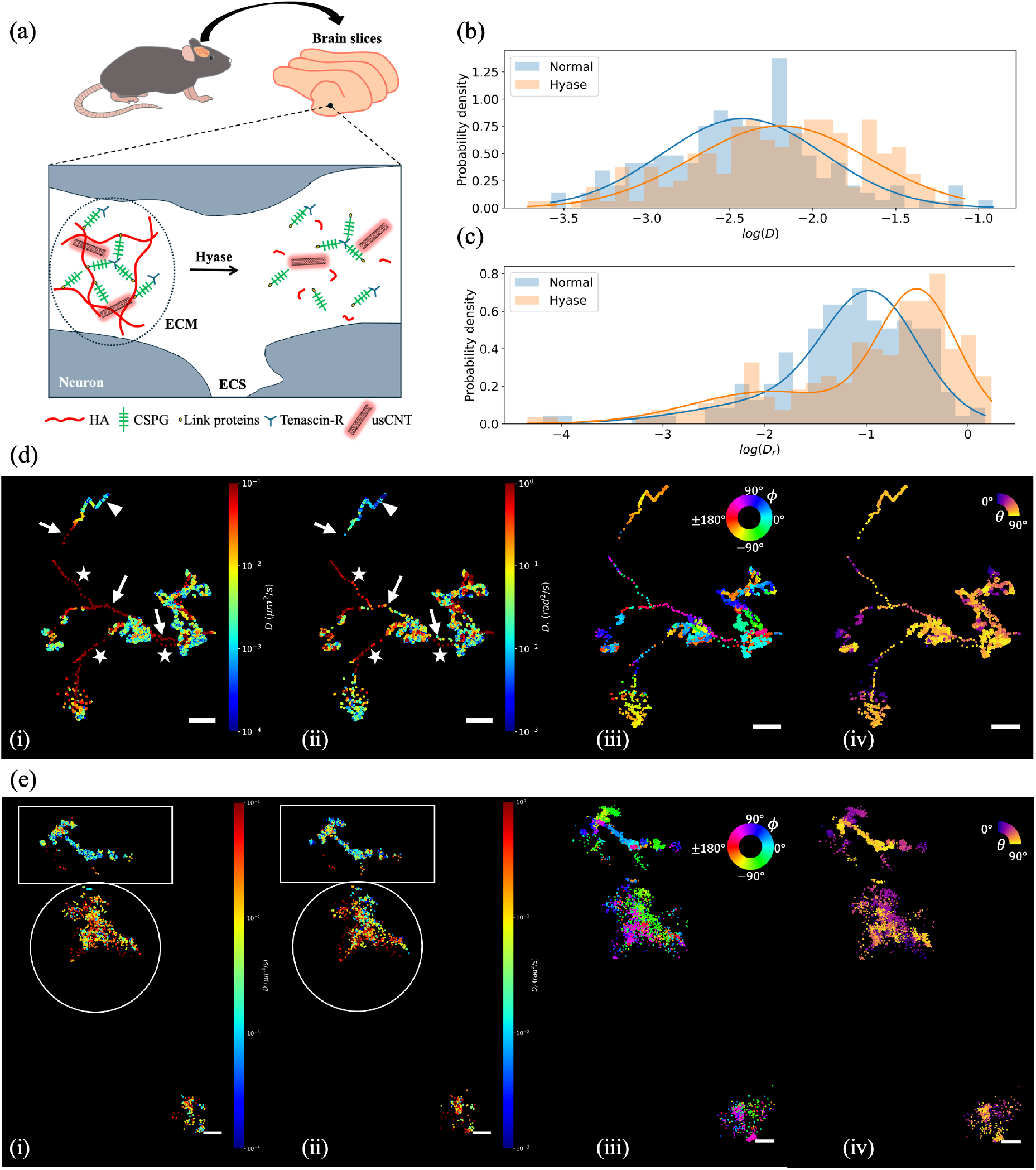
(a) Schematic of the effects of hyaluronidase (Hyase) on the extracellular matrix (ECM) within the brain extracellular space (ECS). (b-c) The probability distributions of log(*D*) (b) and log(*D*_*r*_) (c) are fitted by a Gaussian Mixture Model. We analysed 173 trajectories for normal brain slices and 194 trajectories for Hyase brain slices (Figure S9). (d-e) 2D ECS images of instantaneous translational (i)) and rotational (ii) diffusion and their *ϕ* (iii) and *θ* (iv) map for normal (d) and Hyase (e) brain slices. Scale bar: 0.5 *µ*m.

Figure 6(d) presents typical single uCCNT trajectories color coded by instantaneous translational and rotational diffusion coefficients calculated on a time window of 330 ms thus corresponding to 10 data points, as well as by in plane and out of plane angles. Diffusion in the brain ECS exhibited a strong heterogeneity not seen in homogenous medium nor in agarose gels. Most interestingly, this analysis indicates that local variations of *D* and *D*_*r*_ are not systematically correlated revealing local organization of the brain ECS that translational diffusion analysis alone would not reveal.

More precisely, regions displaying both high *D* and high *D*_*r*_, without clear trajectory or orientation signatures of confinement, are consistent with low effective viscosity and weak structural restriction, as illustrated by the stars in Figure 6(d)(i,ii). Conversely, relatively fast translational diffusion accompanied by reduced *D*_*r*_, persistent nanotube alignment, or constrained trajectory geometry indicates steric or anisotropic confinement, as shown by the arrows in Figure 6(d)(i–iv). In other regions, both *D* and *D*_*r*_ are low and clear confinement signatures are observed, suggesting contributions from both effective viscous resistance and structural obstacles, as illustrated by the triangle in Figure 6(d)(i–iv). Overall, the joint analysis of, trajectory geometry, and orientation maps allows the respective contributions of effective viscosity and structural confinement to be disentangled.

To gain further insights about the impact of brain ECS organization we finally performed a chemical modulation of the ECS structure by a standard enzymatic disruption of the ECM structure. In the intact ECM, hyaluronan (HA) indeed forms a continuous backbone that anchors chondroitin sulfate proteoglycans (CSPGs) and associated crosslinking proteins [76, 77] (Figure 6(a)). However, upon hyaluronidase (Hyase) application, HA can enzymatically cleaved into short fragments, causing destructuration of the ECM. This remodelling of the ECM is expected to reduce structural integrity and affecting both local viscosity and the physical barriers that impede molecular diffusion [26] (Figure 6(a)). Figure 6(b,c) and Figure S8 summarize the translational and rotational diffusion behaviors of uCCNTs in intact and degraded ECM slices. The probability distributions of log(*D*) and log(*D*_*r*_) which can be fitted with a Gaussian Mixture Model, show an increases in both translational and rotational diffusivities. The dominant peak of the translational diffusion coefficient shifts from *D*_Normal_ = 0.004 *µ*m^2^*/*s to *D*_Hyase_ = 0.006 *µ*m^2^*/*s, while the characteristic rotational diffusivity increases from *D*_*r*,Normal_ = 0.1 rad^2^*/*s to *D*_*r*,Hyase_ = 0.3 rad^2^*/*s. Together, these observations suggest that Hyase-mediated ECM disruption globally reduces both ECS viscosity and diffusion obstacle crowding. Notably, a detailed analysis of the spatial correlations between these diffusivities shows no systematic correlation, highlighting further that orientational tracking yields complementary insights that translational tracking alone cannot provide as seen in Figure 6(e)(i-iv) which displays examples of single uCCNT trajectories in enzymatically degraded brain slices. Distinct behaviors can indeed be distinguished, which we attribute to the heterogeneous effect of Hyase. The top region (square) does not appear to have been strongly affected by Hyase displaying rather heterogeneous areas where diffusion can be hindered and slow as revealed in control brain slices. The other bottom regions (circle) display more homogeneous behaviors where uCCNTs display both facilitated translational and rotational diffusion in large domains. Interestingly, in such ‘tortuosity map’, displaying these large domains, a finer analysis of nanotube orientations reveals the presence of distinct subdomains characterized by well-defined uCCNT alignments (Figure 6(e)(iii,iv)). One may attribute this observation to a localized structuration of the ECM, which imposes spatial constraints on uCCNT orientations within confined microenvironments which may involve different components of the ECM, or cellular processes heterogeneity. Due to low viscosity of the medium upon HA digestion (Figure 6(e)(i)), uCCNTs undergo rapid and efficient exploration of these subdomains, also exhibiting fast rotational diffusion as demonstrated by the high in *D*_*r*_ values in the circled area as compared to rectangular one (Figure 6(e)(ii)). As can be seen in Figure 6(e)(iii,iv), uCCNTs are however more likely to move along their long axis, a similar observation as agarose experiment. Therefore in this non-Brownian motion situation imposed by local structure of the ECM, uCCNTs exhibit directional properties which defines these patches while translational diffusion, likely facilitates transitions between these structured regions. Altogether, such ‘tortuosity map’ featuring well-defined patches with stable and distinct orientations could not have been inferred from translational tracking alone. To unambiguously establish the existence of such local ECM organization in degraded brain tissue, further investigations will be required. These would require finer modulation of ECM properties which is beyond the scope of the present study which aimed to illustrate the potential orientational tracking to provide novel rheological understanding of complex tissue.

## Conclusion

In summary, we presented an innovative approach to investigate local rheological features of complex environments at high spatial resolution. This achievement was enabled by the unique properties of fluorescent carbon nanotubes combined with single particle orientation tracking in the SWIR domain. High aspect ratio of CCNTs facilitates orientation measurements at the single-particle level, while their exceptional brightness and photostability in the SWIR range allow for deep and stable imaging of individual particles over extended periods, even in intricate environments. Additionally, the bio-compatibility of CCNTs, together with the ability to tailor their length through quantum defect functionalization, provides a versatile approach: this adaptability allows precise tuning of their rotational behavior to match the dimensions of the environment, making them suitable for capture and dynamic analysis at the single nanotube level. By jointly analyzing trajectory morphology, translational diffusion, and rotational dynamics of individual CCNTs, we could disentangle the respective contributions of effective viscosity and structural confinement in complex environments. We validated this approach in the brain extracellular space, a complex, key yet poorly understood microenvironment whose structural and compositional properties are known to profoundly influence brain function in both physiological and pathological contexts. Using ultrashort CCNTs, we were able to measure the diffusion of single CCNT within the ECS, revealing its structural complexity and role of ECM. This work paves the way for new strategies to explore heterogeneous environments at the nanoscale, with broad applications ranging from biology to materials science.

## Methods

### Radially and azimuthally polarized (raPol) microscope

A dichroic mirror (DM1, DMLP900R) reflects a circularly polarized laser to excite the sample above the objective lens (OL, Nikon CFI75Apo25XCW). The emitted fluorescence is collected by the objective, filtered by DM1 and a longpass filter (LP, Chroma, ET900LP) in order to eliminate completely the excited laser light. It is then focussed by a tube lens (TL, f=200 mm) to form an intermediate image. A second dichroic mirror is added (DM2), similar to DM1 but oriented in a perpendicular plane to compensate the small elliptical polarization effect added by DM1. A 4f system (L1,f1=100 mm; L2, f2=200 mm) is used to engineer the fluorescence at the back focal plane (BFP). The vortex half waveplate (VWP,Thorlabs WPV10L-980) is inserted at the BFP to convert the radially and azimuthally polarization into x- and y-polarization, respectively. A polarizing beamsplitter (PBS, Thorlabs CCM1-PBS225/M) is placed after L2 to separate the x-radial and y-azimuthal polarized fluorescence. Two mirror (M1,2) and a prism mirror (PM, Thorlabs MRAK25-P01) are used to generate side by side two images on a single InGaAs camera (Ninox 640, Raptor).

### Polarization and phase diversities

In standard phase retrieval algorithms, which do not account for polarization, a Z-stack, i.e. a series of intensity images captured at different focal distances from the optimal focus, can provide the required phase diversity. This diversity not only helps the algorithm avoiding local minima during convergence but also enables discrimination between vortices with opposite topological charges. However, for BDPP determination which encodes the polarization information, a Z-stack alone cannot differentiate the true *J* from its unitary transformations *J*_*U*_ = *U*·*J*, where *U* is a constant unitary matrix (*UU*^*†*^ = *I*), which produces identical intensity patterns across the stack. Therefore, polarization diversity must also be introduced which is performed by placing a *λ/*4 placed after the VWP (Figure 2(a)).Rotating the *λ/*4 plate indeed generates multiple polarization projections of the output, providing additional constraints for the vectorial phase retrieval. In practice, the phase diversity is obtained by a Z-stack made of images acquired between -2 and 2 mm around the reference focal plane by increments of 400nm, and the polarization diversity by rotated the *λ/*4 waveplate from 0 to *π/*2 in steps of *π/*6. Each diversity DSF was acquired by averaging 20 images with 100ms integration time.

### CCNT preparation

CCNTs were prepared following the protocol reported in previous work [43, 78]. Briefly, monochiral (6,5)-SWCNTs were sorted by aqueous two-phase extraction (ATPE) as described previously [79] and shortened by extended tip sonication. They were then functionalization with luminescent oxygen defects via a Fenton-like reaction with copper(II) sulfate (CuSO_4_(H_2_O)_5_) and sodium L-ascorbate according to the established protocol [78] and eventually followed by surfactant exchange to PL-PEG5000 (Avanti Lipids) via dialysis for biological applications.

### Spin-coating uCCNT Samples

The DOC-uCCNTs solution was diluted 1:9 with distilled water. A coverslip is covered by 70 *µ*L polyvinypyrrolidone (PVP) 3 times. A 30 *µ*L aliquot of the diluted solution of uCCNTs was deposited onto the coverslip, which was then spin-coated at 3000 rpm for 30 s to immobilize the uCCNTs on its surface. Then, It is imaged by SPoT microscope with 100ms expsure time.

### 5% agarose-uCCNT Samples

A 5% agarose solution was prepared by dissolving 0.75 g of agarose in 15 mL of distilled water. Subsequently, 0.5 *µ*L of DOC sp^3^ uCCNT solution was mixed with 250 *µ*L of 5% agarose. The mixture was deposited onto a coverslip and allowed to sit for 30 min at room temperature to ensure uCCNTs immobilization.

### Single particle orientation tracking (SPoT)

The 2D trajectory is reconstructed by single particle tracking (SPT) algorithm. Multi-particle tracking can be performed using the trackpy library, which is based on the Crocker– Grier algorithm [80]. The algorithm formulates trajectory linking as a linear assignment problem, where candidate particle displacements are optimized globally to maintain reliable particle identities over time. This enables robust tracking in challenging conditions, including high particle density, closely spaced localizations, intermittent blinking, and occasional missing detections. Since the 3D orientation at each time point is now available, the same temporal axis can be used to construct the corresponding orientational trajectory. Subsequently, we perform an angle unwrapping procedure to obtain the continuouse orientation trajectory.

### Glycerol-CCNT Samples

To prepare glycerol solutions, 90% glycerol was obtained by mixing 9 mL of glycerol with 1 mL of distilled water. Similarly, 80% and 70% glycerol solutions were prepared by mixing appropriate volumes of glycerol and distilled water. Subsequently, 0.5 *µ*L of DOC sp^3^ long CCNT solution was added to 250 *µ*L of each glycerol solution (90%, 80%, and 70%, respectively). Samples were mounted between a glass slide and a coverslip and incubated for 30 min to minimize flow prior to SPoT measurements..

### 2% agarose-CCNT Samples

A 2% agarose solution was prepared by dissolving 0.6 g of agarose in 15 mL of distilled water. Subsequently, 0.5 *µ*L of the DOC sp^3^ long CCNT solution was mixed with 250 *µ*L of 2% agarose. The mixture was then deposited between a glass slide and a coverslip and allowed to solidify for 30 min at room temperature.

### Imaging immobilized uCCNTs through acute mice brain tissue slices

DOC-uCCNTs were immobilized on a coverslip by spincoating. Mice (strain C57BL6/JRj purchased from Janvier Laboratories and reared at the University of Bordeaux animal facilities) were euthanized by cervical dislocation, and whole brains were swiftly extracted. Acute brain slices (60−100 *µ*m thick) were prepared in an ice-cold, carbogenated sucrose-based cutting solution containing (in mM): 210 sucrose, 2.5 KCl, 26 NaHCO_3_, 1.25 NaH_2_PO_4_, 10 D-glucose, 4 MgSO_4_, and 0.5 CaCl_2_. The solution was continuously equilibrated with 95% O_2_/5% CO_2_ to maintain a pH of 7.3–7.4 and an osmolarity of approximately 300–320 mOsm. Brains were rapidly removed and immersed in the ice-cold cutting solution, and coronal slices were prepared using a vibratome in the same solution. Recordings were performed in the same solution equilibrated at RT. Experiments were performed following the European Union directive (2010/63/EU) on protecting animals used for scientific purposes. Brain slices were placed on a coverslip containing immobilized uCCNTs, and the probes were imaged through the tissue slices by raPol microscope. A total of 200 frames were recorded for each uCCNT.

### Organotypic Slice

Organotypic brain slice cultures were prepared as previously described [25, 28]. Hippocampal slices were obtained from postnatal day 5 to day 7 Sprague–Dawley rat pups and cultured for up to 14 days on hydrophilic polytetrafluoroethylene (FHLC) membranes (0.45 *µ*m, Millipore) placed on Millicell cell culture inserts (Millipore, 0.4 mm thickness, 30 mm diameter). The culture medium was changed every 2–3 days. For single-particle tracking experiments, uCCNTs were prepared as previously described [43]. Slices were incubated in 190 *µ*L of culture medium mixed with 10 *µ*L of uCCNT solution for 2h at 35 ^◦^C under 5% CO_2_. Imaging was performed in HEPES-based artificial cerebrospinal fluid (aCSF) containing 145 mM NaCl, 4 mM KCl, 2 mM CaCl_2_·2H_2_O, 1.0 mM MgCl_2_·6H_2_O, 10 mM HEPES, and 10 mM D-glucose (osmolarity ∼ 310 mOsm, pH 7.35), with the temperature maintained at 35 ^◦^C using a temperature controller (Tokai Hit). The slices was imaged through a coverslip.

### Drift correction

Frame-to-frame displacements (*dx, dy*) were computed for each molecule. The average displacement per frame was then used to estimate the global drift of the system. To reduce noise, the drift signal was smoothed using a rolling window (window = 5 frames). The cumulative drift over time was fitting by linear function, producing fitted drift components. These fitted components represent the systematic drift of the system. To minimize the influence of outliers, molecule positions that deviated by more than three standard deviations from their mean positions were excluded from the analysis. Finally, the estimated drift was subtracted from the raw trajectories to obtain corrected molecule positions.

### Hyaluronidase treatment

Hippocampal slices were incubated with hyaluronidase enzyme (H3631, Sigma) at a dilution of 50 U*/µ*L for 4 h, as previously described [25]. Subsequently, the slices were incubated with uCCNTs for 2h and imaged in HEPES-based artificial cerebrospinal fluid (aCSF).

## Acknowlegments

We thank the Cell Biology Facility for managing cell biology-related activities, especially Delphine Bouchet. We also thank Xavier Le Guevel (Université Grenoble-Alpes) and Andreas Reisch (Université de Strasbourg) for providing the polymeric nanoparticles (GNPs). L.C. and L.G. acknowledges financial support from the European Research Council Synergy grant (951294). L.C. acknowledges support from Agence Nationale de la Recherche (EUR Light&T, PIA3 Program, ANR-17-EURE-0027), the France-Bioimaging National Infrastructure (ANR-10-INBS-04-01), and the Idex Bordeaux (Grand Research Program GPR LIGHT). LR is financially supported by the China Scholarship Council (Grant No. 202208330010). L.A.A-C. and S.B. acknowledge support from Agence Nationale de la Recherche (ANR-21-CE24-0014). F.L.S.,and J.Z. acknowledge funding from the European Research Council (ERC) under the European Union’s Horizon 2020 research and innovation programme (Grant Agreement No. 817494 “TRIFECTS”). J.Z. acknowledges additional support by the Deutsche Forschungsgemeinschaft (DFG, German Research Foundation) under Germany’s Excellence Strategy for the Excellence Cluster “3D Matter Made to Order” (EXC-2082/1 - 390761711).

## Data availability statements

The data that support the findings of this study are openly available in Zenodo at https://doi.org/10.5281/zenodo.20272734.

