## Supplemental information for "Nanoscale rheological heterogeneity revealed by Single Particle orientation Tracking (SPoT) of ultrashort carbon nanotubes in brain tissue"

### A Derivation of the mean angular standard deviation

The orientation of a dipole is represented by the unit vector

$$\vec{\mu}(\theta, \phi) = [\sin\theta\cos\phi,\ \sin\theta\sin\phi,\ \cos\theta]^T, \tag{S1}$$

where $\theta$ and $\phi$ denote the polar and azimuthal angles, respectively.

The infinitesimal line element on the sphere is

$$dS^2 = d\theta^2 + \sin^2\theta d\phi^2, \tag{S2}$$

which gives the local metric of the spherical surface. Consequently, the infinitesimal surface area is

$$dA = \sin\theta d\theta d\phi. \tag{S3}$$

Assuming that the uncertainties in the polar and azimuthal directions are sufficiently small, the uncertainty region can be approximated by a rectangular patch on the tangent plane. Its area is therefore

$$A \approx \sin\theta\sigma_\theta\sigma_\phi, \tag{S4}$$

where $\sigma_\theta$ and $\sigma_\phi$ are the standard deviations of the estimated $\theta$ and $\phi$.

To describe this uncertainty using a single angular quantity, we define an equivalent spherical cap whose projected area equals that of the uncertainty patch. The patch area is therefore equated to the area of a circle with radius $r$ on the tangent plane,

$$A = \pi r^2, \tag{S5}$$

yielding

$$r = \sqrt{\frac{\sin\theta\sigma_\theta\sigma_\phi}{\pi}}. \tag{S6}$$

For a unit sphere, the tangent-plane radius r is related to the corresponding angle $\sigma_\delta$ through the geometric relation

$$r = \sin(\frac{\sigma_\delta}{2}), \tag{S7}$$

which leads to

$$\sigma_\delta = 2\arcsin\sqrt{\frac{\sin\theta\sigma_\theta\sigma_\phi}{\pi}}. \tag{S8}$$

The quantity $\sigma_\delta$ represents the angle of an equivalent circular uncertainty region on the unit sphere and provides a single scalar measure of orientation precision by combining the estimation uncertainties in both the polar and azimuthal directions.

#### B Supplemental Figures

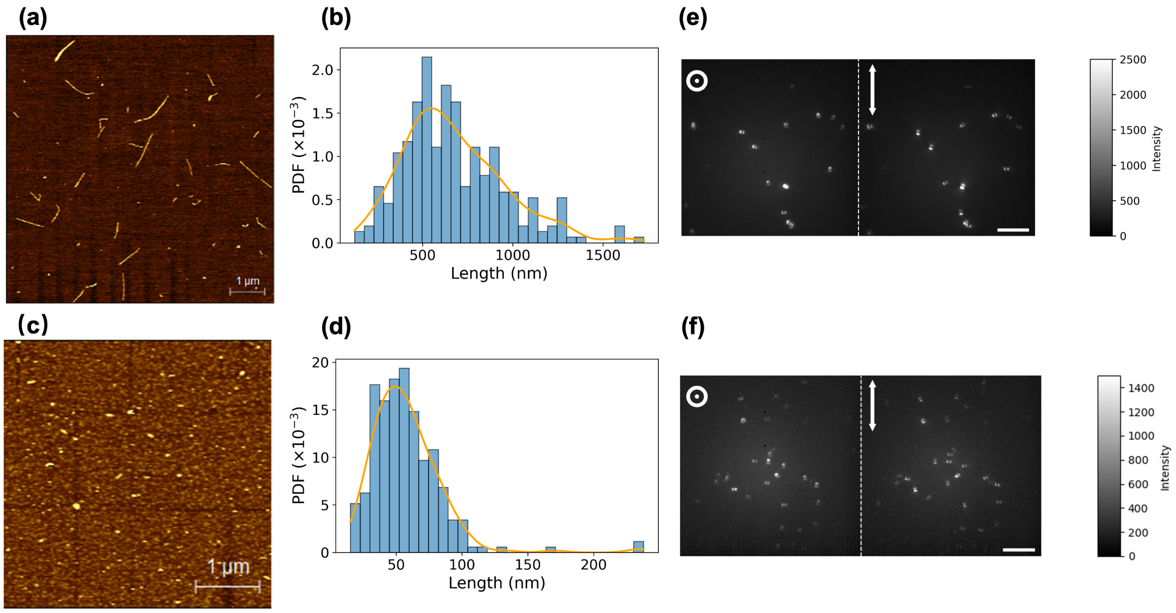

Figure S1: **AFM images and distribution of length of CCNTs and ultrashort CCNTs.** (a,c) AFM images of CCNTs (a) and ultrashort CCNTs (c). (b,d) Corresponding length distributions of 288 CCNTs (b) and 237 ultrashort CCNTs (d). The length were analyzed by using Gwyddion software. (e,f) raPol microscope images of CCNTs (e) and ultrashort CCNTs (f). Similar DSF patterns are observed for both lengths of nanotubes. The only difference is that CCNTs display sliglthy brighter signal. Left: X-radial images. Right: Y-azimuthal images. Scale bar: 10 $\mu$m.

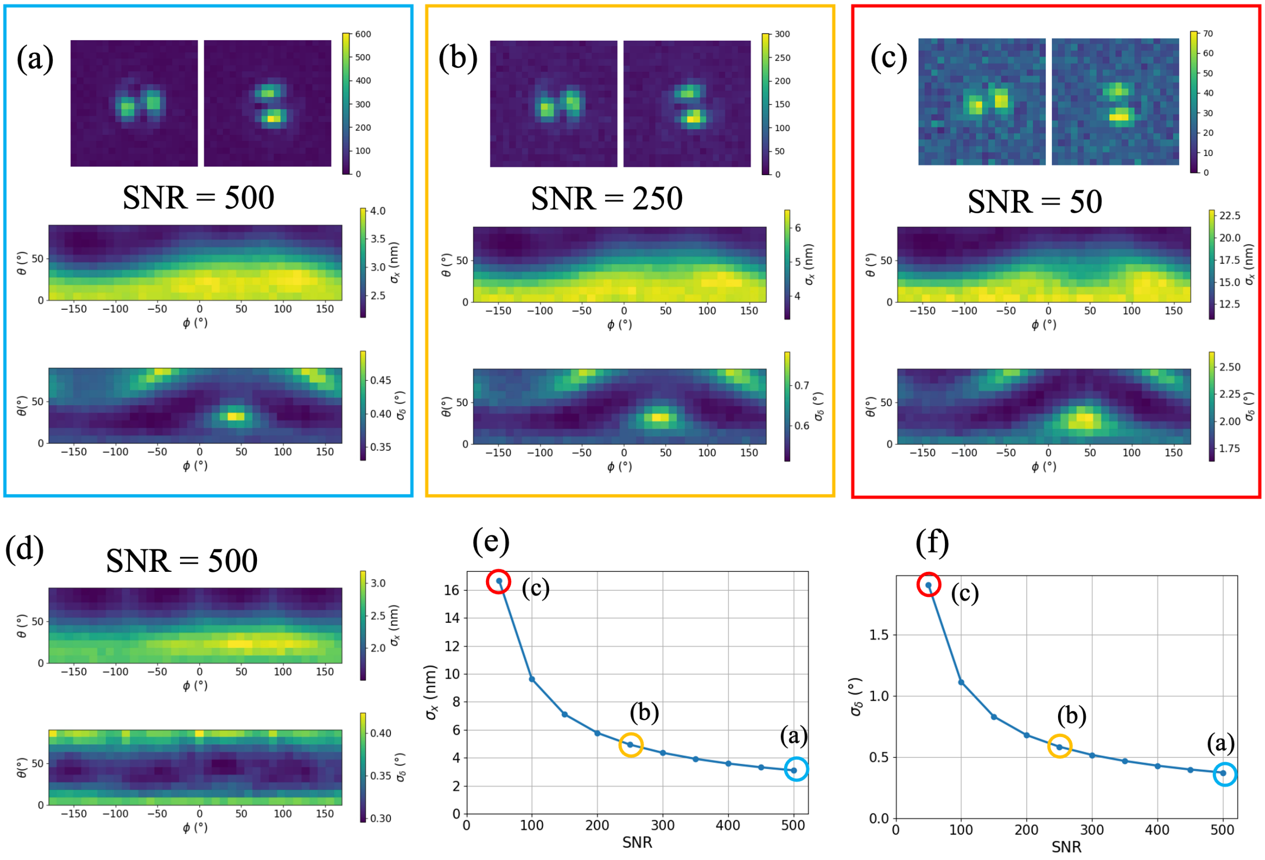

Figure S2: **CRLBs calculated for localization ($\sigma_r$) and orientation ($\sigma_\delta$) of single dipoles detected by the raPol microscope.** (a-c) Simulated DSFs ($\theta = 90°, \phi = 0°$) and the corresponding orientation-dependent CRLBs of single dipoles at different SNRs using experimental BDPP which contains experimental abberations. (d) Orientation-dependent CRLBs of single dipoles using simulated BDPP with resulting values of $\sigma_\delta = 0.3$, $\sigma_r = 2{,}3$. (e,f) Orientation-averaged CRLBs for localization (e) and orientation (f) as a function of SNR.

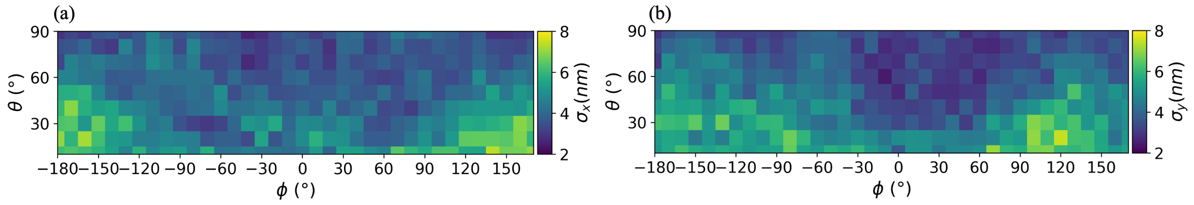

Figure S3: **Deep-SMOLM estimation precision of localization.** (a) Precision of x ($\sigma_x$). (b) Precision of y ($\sigma_y$).

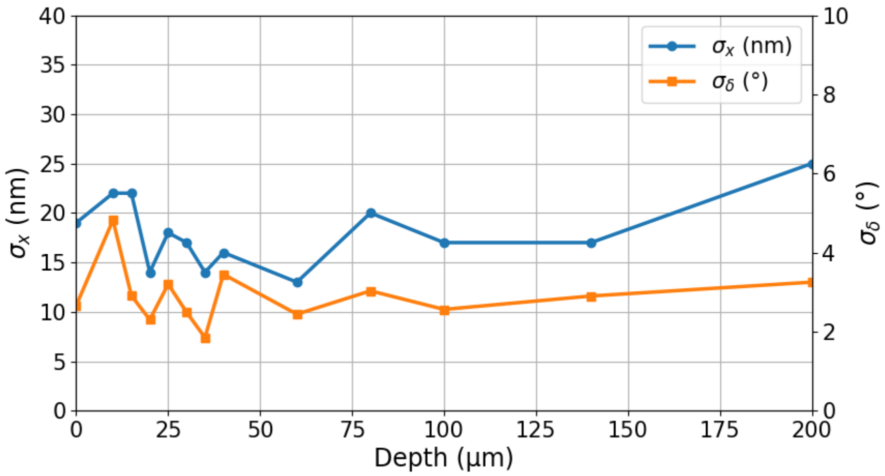

Figure S4: **Depth dependance of localization and orientation precision in agarose gel**

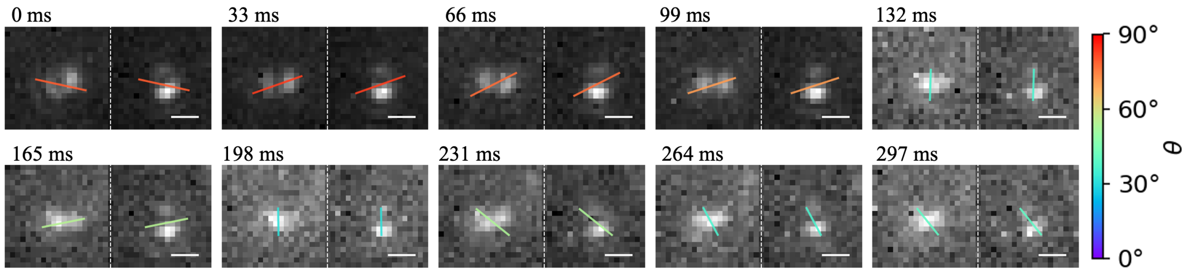

Figure S5: **Typical DSF measurements for CCNTs diffusing in 90% glycerol solutions.** Color bar: $\theta$. Left: X-radial DSFs. Right: Y-azimuthal DSFs. Scale bar: 1 $\mu$m.

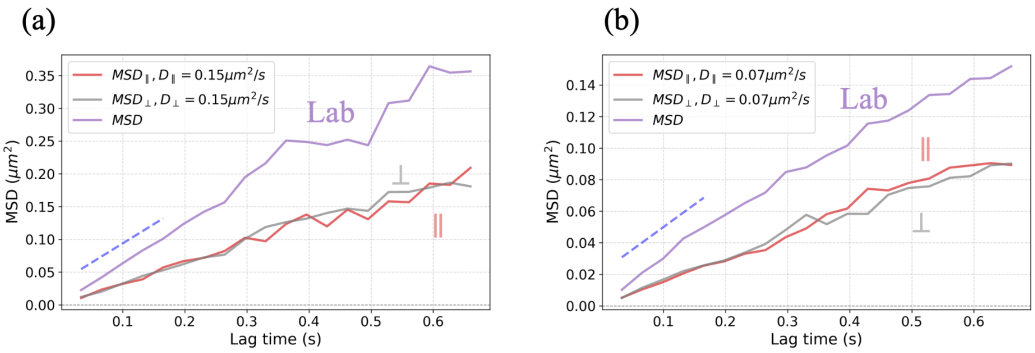

Figure S6: **MSDs of CCNTs trajectories measured in glycerol and calculated in the lab frame (MSD) and in the nanotube frame (MSD$_{\parallel/\perp}$** (a) 70% glycerol. (b) 80% glycerol. Blue dashed line: the slope is $2(D_\parallel + D_\perp)$.

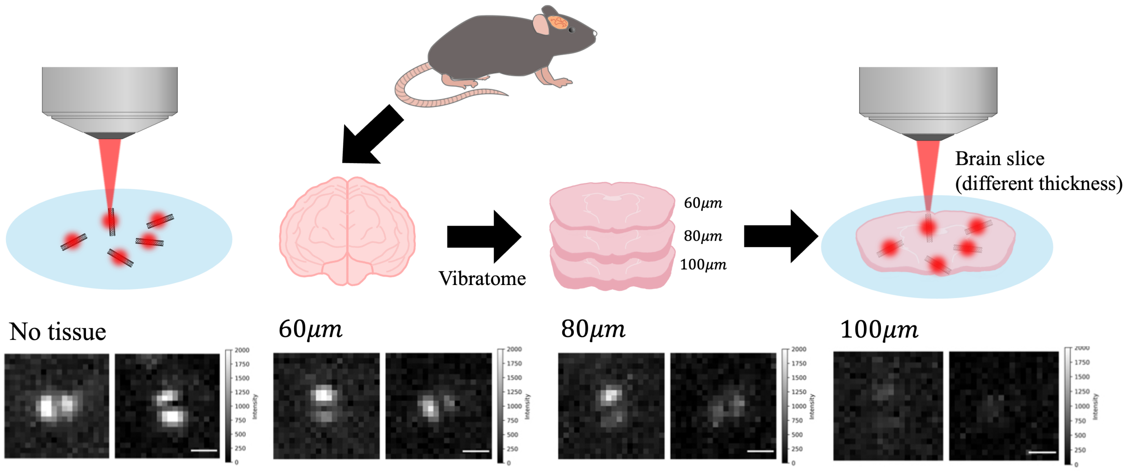

Figure S7: **Depth dependance of localization and orientation precision in acute brain slices.**

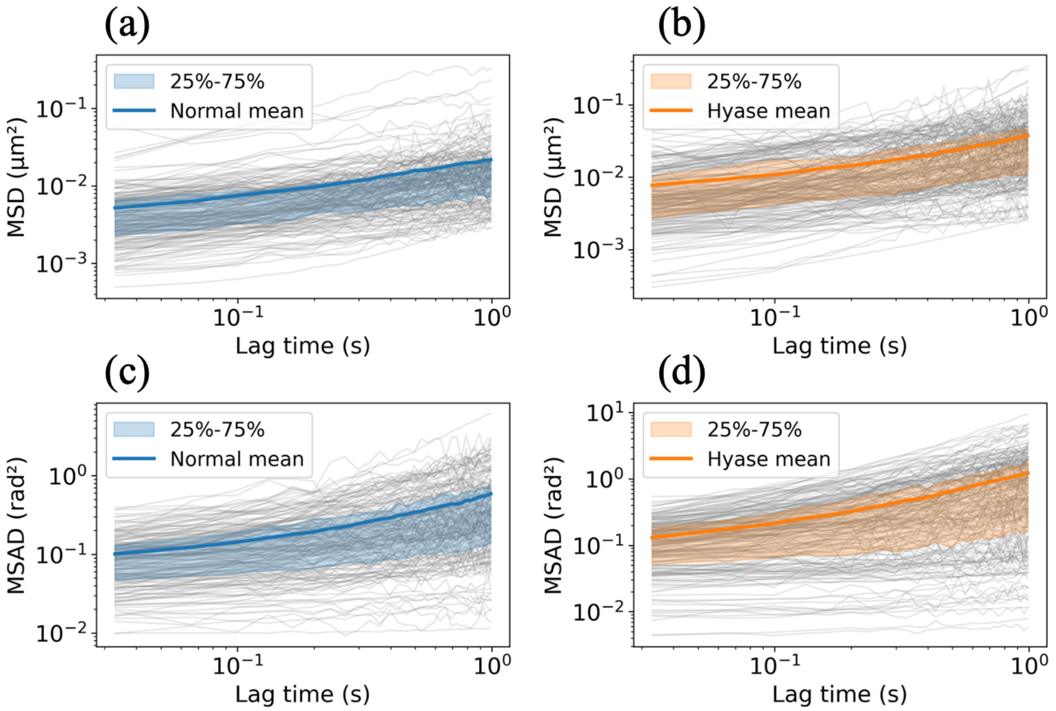

Figure S8: **MSD and MSAD analysis for uCCNTs diffusing in brain slices.** Single trajectories (grey lines) and time-ensemble-averaged (color lines) MSDs (a,b) and MSADs (c,d). Blue: Normal brain slices. Orange: Hyase brain slices. Shaded area: percentile.

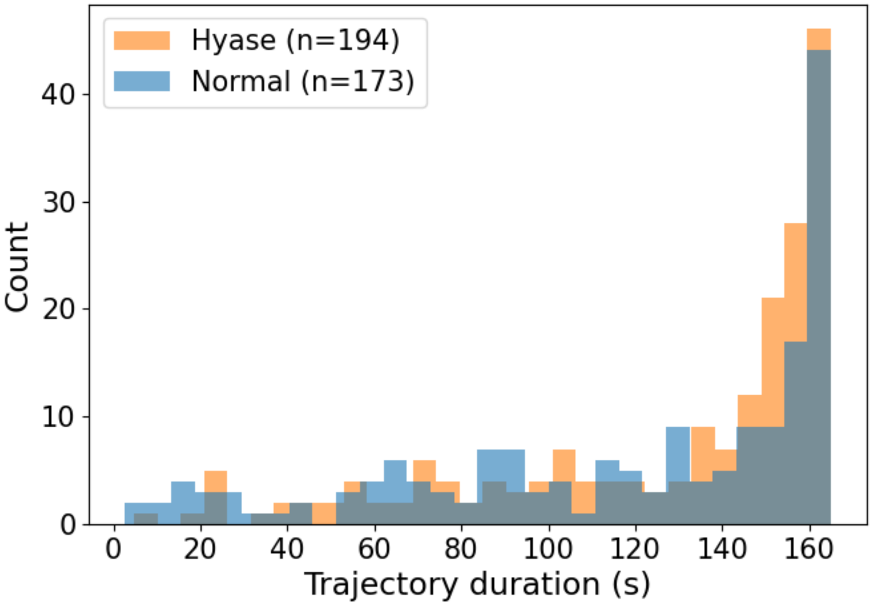

Figure S9: **The trajectory duration distribution. n: the number of trajectories.**
